# Complexome profile of mitochondrial complexes in the myzozoan parasite of oysters, *Perkinsus marinus*

**DOI:** 10.64898/2025.12.18.695115

**Authors:** Andrew E. Maclean, Orsola Iorillo, Drahomíra Faktorová, Monika Singh, Suzanne McGill, Julius Lukeš, Lilach Sheiner

## Abstract

Mitochondrial complexes, such as the mitochondrial electron transport chain (mETC), the F_1_F_o_-ATP synthase and the mitochondrial ribosome, are centrally important for mitochondrial function. Recently, an unexpected diversity in the composition of these complexes across unicellular eukaryotic lineages, with the apicomplexan parasites featuring prominently, has been revealed. However, whether the observed enlarged and divergent mitochondrial complexes are conserved across the wider Myzozoan lineage has not been investigated. Here, using a complexome profiling proteomic approach, we have characterised the composition of the mitochondrial complexes of the myzozoan parasite of oysters, *Perkinsus marinus*. We show that it shares with *Plasmodium* and *Toxoplasma* huge ATP synthase and mETC complexes. Moreover, the *Perkinsus* mitoribosome possesses many of the divergent features recently discovered in apicomplexans, such as an expanded subunit repertoire and the inclusion of RAP-domain and ApiAP2-like proteins as ribosomal proteins. Our study reveals the divergent subunit composition of mitochondrial complexes is an ancestral and highly conserved feature of Myzozoa.

## Introduction

Mitochondria are double membrane bound organelles which are a unique feature of eukaryotes and, despite reduction or loss in some lineages, play an ever-increasing number of vital roles in their biology, ranging from energy metabolism to biosynthesis of key metabolites [1]. One key mitochondrial function is oxidative phosphorylation, which requires the coordinated function of several large mitochondrial complexes. The mitochondrial electron transport chain (mETC) is a series of complexes embedded in the inner mitochondrial membrane (IMM), named complex I (NADH dehydrogenase), complex II (succinate dehydrogenase), complex III (cytochrome *bc*_1_ complex) and complex IV (cytochrome *c* oxidase). These complexes are required to transfer electrons harvested from a variety of metabolic pathways to oxygen. This process is coupled with proton translocation across the inner mitochondrial membrane, creating an electrochemical gradient which is utilised by complex V (F_1_F_o_-ATP synthase) to generate ATP [2]. While most subunits of the mETC are encoded in the nucleus and imported into the organelle, several key subunits are encoded by the mitochondrial genome. They therefore require another mitochondrial complex, the mitochondrial ribosome (mitoribosome), for their translation. Mitochondrial complexes thus enable many core functions of the organelle.

Historically, most mitochondrial research was focused on oxidative phosphorylation and mitochondrial translation in a small group of model organisms that are closely related to one another, primarily mammals and yeast, resulting in an incomplete view of mitochondrial biology. However, recent research, particularly in unicellular eukaryotes, has started revealing that mitochondrial biology displays considerable diversity [3].

The Myzozoa are a group of unicellular eukaryotes which contains both parasitic and free-living species [4,5]. Parasitic lineages include species that have a huge impact on global health, the economy and the environment. For example, the phylum Apicomplexa brings together parasites that cause global diseases of both humans and livestock, most notably the malaria and toxoplasmosis caused by *Plasmodium* spp. and *Toxoplasma gondii*, respectively. Another example are the perkinsids, marine parasites that infect molluscs, including oysters, causing significant economic and environmental impact [6]. Myzozoans also include lineages that contain free-living algae and coral symbionts, such as the chrompodellids and dinoflagellates [4].

Recent studies showed that the apicomplexans display mETC and F_1_F_o_-ATP synthase complexes that are highly divergent from previously studied systems, both in composition and structure. All members have lost complex I, and there have been multiple parallel losses of mETC complexes in numerous lineages [7,8]. Of those that retain mETC complexes, including *Plasmodium* and *Toxoplasma*, the individual complexes have inflated subunit compositions compared to canonical complexes and display unique structures [9–16]. Importantly, a similar pattern is seen for F_1_F_o_-ATP synthase [10,11,17–19]. These new subunits correlate with novel features and functions of the corresponding complexes. For example, clade-specific subunits mediate ATP synthase hexamerization which is important for cristae formation [17], and mETC supercomplex formation [9]. Likewise, apicomplexan mitoribosomes are substantially distinct from their homologs in other eukaryotes. While apicomplexan mitochondrial genomes encode the lowest number of protein-coding genes known [20], their ribosomal RNAs (rRNAs) are uniquely extremely fragmented [21,22]. In *Toxoplasma* numerous clade-specific proteins take place in mitigating the extreme rRNA fragmentation, many of which seem conserved across the Myzozoa [23,24].

Through proteomic and structural investigations, we now have a much clearer understanding of apicomplexan mitochondrial complexes. However, we know close to nothing about the other myzozoan lineages. Intriguingly, many of the apicomplexan-specific subunits have putative homologs in these other lineages, hinting at clade-wide conservation. To gain a fuller picture of myzozoan mitochondrial biology, here we investigate mitochondrial complexes in a non-apicomplexan group. We chose to examine the marine parasite *Perkinsus marinus*, the best-studied member of the perkinsids, which is a sister clade to the dinoflagellates [4]. Perkinsids diverged close to the split with the clade containing apicomplexans and chrompodellids, making it a suitable sampling point for myzozoan diversity. Additionally, recent genetic and cell biology studies [25,26] make *P. marinus* a tractable model that will enable answering questions emerging from this work.

Here, we applied proteomic complexome profiling to *P. marinus* in order to gain a broad view of the protein composition of its mitochondrial complexes and understand the conservation of highly divergent organellar features across myzozoans.

## Results

### Isolation of *Perkinsus* mitochondria

Complexome profiling has proven a powerful proteomic technique for uncovering the composition of organellar protein complexes [27–29]. To study the mitochondrial complexes of *P. marinus*, we decided to take a dual approach, performing complexome profiling on both mitochondrially-enriched and whole-cell material.

To isolate material enriched in mitochondria, *P. marinus* cells (strain ATCC 50983) were lysed *via* mechanical disruption using glass beads, followed by differential centrifugation to enrich organellar material (see Materials and Methods). Immunoblot analysis of obtained fractions, using antibodies against the conserved ATP synthase beta subunit (ATPβ) (Fig 1A), confirmed mitochondrial enrichment. Antibodies against both a universally conserved cytosolic glyceraldehyde 3-phosphate dehydrogenase (GAPDH) and a nuclear histone H3 showed a strong signal in the total fraction, but absence or a decreased intensity in the mitochondrial fraction (Fig 1A), in support of the removal of these cell compartments during purification. On the other hand, an endoplasmic reticulum (ER) protein, the chaperone BiP, was also present in the mitochondrial fraction, indicating co-purification with other organelles. These data suggest that the purification protocol was able to obtain material enriched in mitochondrial proteins.

**Figure 1:**
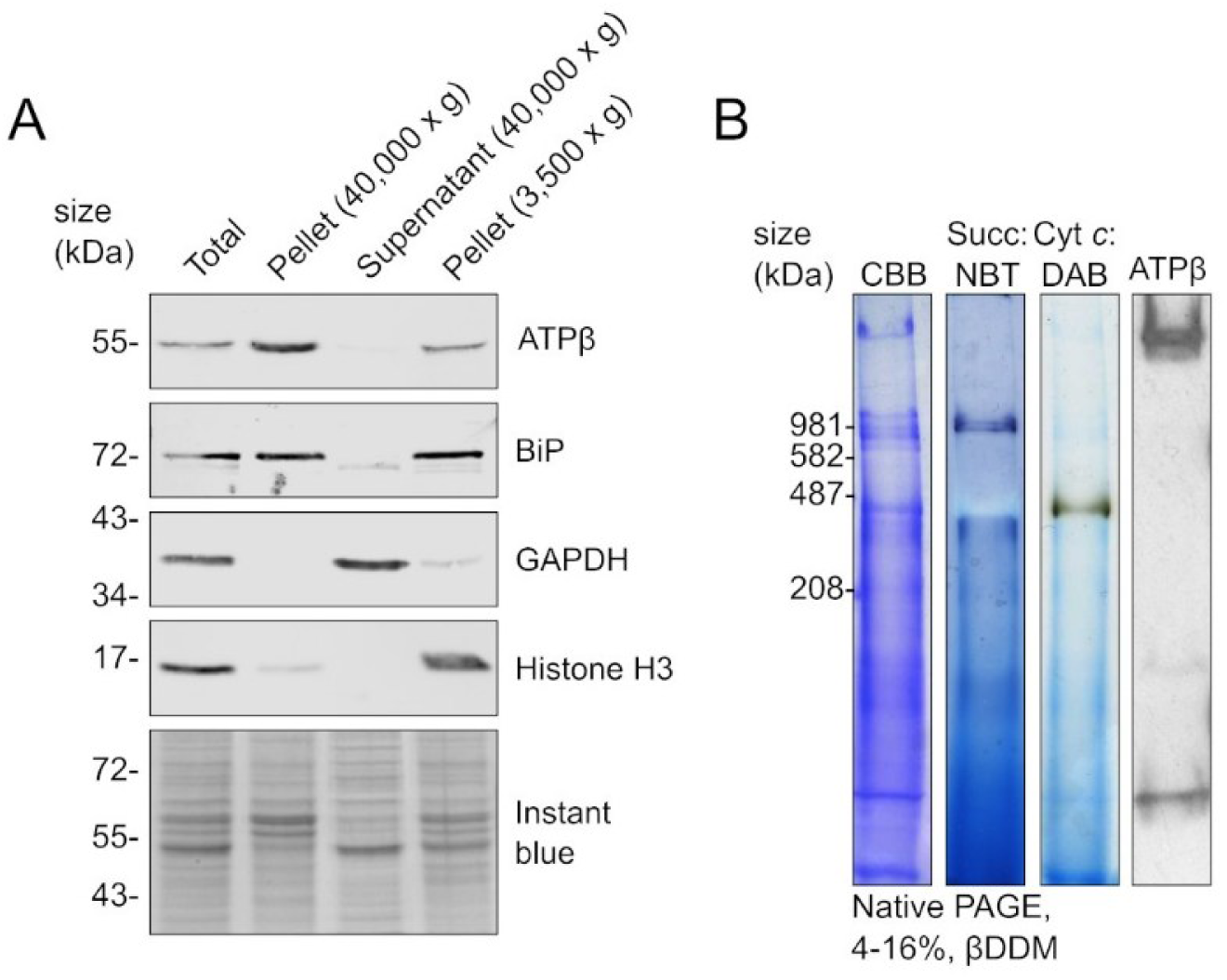
Isolation of mitochondrially-enriched fractions. (A) Immunoblot analysis of total cell lysate (Total) and purification fractions using antibodies against marker proteins of various cell compartments. The pellet from the 3,500 x *g* centrifugation step contains unbroken cells and heavy cell components, from the 40,000 x *g* step the supernatant contains soluble cytosolic proteins, and the pellet mitochondrial and organellar material. 7.5 μg of protein was loaded per lane with equal loading confirmed by instant blue staining (bottom panel). The following antibodies were used: ATPβ (mitochondrion), BiP (ER), GAPDH (cytosol) and Histone H3 (nucleus). (B) Blue-native PAGE analysis of mitochondrially-enriched material treated with βDDM and separated on a 4-16% Bis-Tris gel. Total protein was visualized by Coomassie staining, complex II activity by an in-gel succinate: nitroblue tetrazolium (NBT) activity stain, complex IV activity by an in-gel cyt *c*: 3,3′-Diaminobenzidine (DAB) activity stain and ATP synthase by immunoblot analysis using the anti-ATPβ antibody.

Next, to test whether we have isolated mitochondrial material suitable for complexome profiling, where intact complexes are required, we performed native-PAGE analysis. The mitochondrial material was solubilised in the non-ionic detergent n-dodecyl β-D-maltoside (βDDM) and complexes were separated by size *via* blue-native polyacrylamide gel electrophoresis (BN-PAGE). Coomassie staining of the gel showed numerous, clear, high-molecular-weight bands representing large macromolecular complexes (Fig 1B). To screen for known mitochondrial complexes, we first performed in-gel histochemical stains to test for the activity of mETC complex II and complex IV. By applying the complexes II- and IV-specific stains, we observed activity bands at ∼900 kDa and ∼450 kDa, respectively (Fig 1B), suggesting that in *P*. *marinus* these complexes are significantly larger than those seen in mammalian or yeast systems and similar in size to those described from the best-studied apicomplexans, *Plasmodium* and *Toxoplasma* [10,11]. We next performed immunoblot analysis with the ATPβ antibody to detect the F_1_F_o_-ATP synthase complex. We observed a high molecular weight band above 1 MDa (Fig 1B), suggesting a large complex of similar size to that seen in the apicomplexan parasites [10,17,30]. Additional low-molecular-weight bands were observed, perhaps reflecting monomeric ATPβ or assembly intermediates. Thus, BN-PAGE analysis indicates that the mitochondrially-enriched samples are suitable for further complexome analysis and suggests that *Perkinsus* possesses oxidative phosphorylation complexes that are divergent from commonly studied models and potentially similar to other myzozoans.

To investigate the composition of protein complexes within the whole-cell, *P. marinus* cells (strain PRA-240 (CB5D4;[31]) were lysed in a buffer containing the detergent IGEPAL *via* mechanical disruption with glass beads and the whole-cell protein content was separated in a BN-PAGE gel for complexome analysis.

### Complexome profiling of *Perkinsus* mitochondrial complexes

We then proceeded to perform complexome analysis on both the mitochondrially-enriched material and the whole-cell lysate. The mitochondrially-enriched fraction was treated with 1.7% βDDM to solubilise membrane complexes, which were then separated by BN-PAGE, stained with Coomassie, and the gel lane was cut into 60 gel slices (S1A Fig). Each gel slice was then analysed by mass spectrometry and proteins identified using the Uniprot UP000007800 proteome annotation, supplemented with the sequences of mitochondrially-encoded proteins [25]. Abundance data for every protein for each of the remaining 60 gel slices was calculated using the iBAQ method, before normalisation and hierarchical clustering was performed to generate the complexome profile [32,33]. Two independent complexome profiles, performed with separate mitochondrial isolations, were generated (S1 Table). Between the two replicates, 4,243 individual proteins were detected, with 2,835 common to both datasets (S1B Fig; S1 Table). Further analysis was performed only on proteins detected in both datasets, unless stated otherwise. Two sets of molecular weight markers were run in parallel with the *Perkinsus* material: one composed of soluble proteins and the other containing bovine mitochondrial membrane complexes (S1A Fig). This allowed mass calibration based on the migration of both soluble and membrane-embedded proteins (S1A,C Fig; S3 Table).

For the whole-cell sample, proteins were solubilised in 0.1% IGEPAL, a milder non-ionic detergent, in order to detect weaker inter-complex interactions. Protein complexes were separated by BN-PAGE and the gel was excised into 48 slices (S1A Fig). Samples were analysed as above, with 3,552 proteins detected (S1B Fig; S2 Table). Between the three datasets 1,747 proteins were detected in common (S1B Fig; S1, S2 Tables), with 1,331 detected only in the whole-cell dataset, indicating differential cell component enrichment between the two sample types, as would be expected. A ladder composed of soluble proteins was run in parallel, allowing mass calibration (S1A,C Fig; S3 Table).

Next, to assess whether our complexome datasets had good coverage of the mitochondrial proteome we compared it to organellar datasets from other myzozoan species. We first compared our two mitochondrially-enriched complexome datasets to the hyperLOPIT spatial proteomics data from *Toxoplasma* [34]. Our datasets contained putative *Perkinsus* homologs of between 64-67% of the *Toxoplasma* proteins found in the mitochondrial clusters. This includes 67-70% of proteins belonging to the “mitochondrial membranes” cluster. Our whole-cell sample contained putative homologs of 46% of proteins in the mitochondrial clusters. Finally, comparisons to a curated mitochondrial proteome from *Plasmodium falciparum* [35] showed our mitochondrially-enriched complexome datasets contained homologs of between 57-60% of *Plasmodium* mitochondrial proteins, while the whole-cell dataset contained 38% of them. Overall, these comparisons suggest our complexome profiles have sufficient coverage of mitochondrial proteins, suitable for assessing mitochondrial complex composition.

### Mitochondrial electron transport chain complexes

The multi-subunit complexes of the mETC show considerable compositional diversity across eukaryotes, and recent studies in apicomplexans suggest a much-expanded subunit repertoire compared to the commonly studied opisthokont models [2,3]. In order to determine whether this expanded subunit repertoire extends to other myzozoans, we interrogated our *Perkinsus* complexome datasets for mETC complexes.

We first searched for complex II subunits. While canonically a 4-subunit complex in the bacterial ancestor and in well-studied models such as yeast and mammals, other lineages contain substantially enlarged complexes. For example, plants possess an 8-subunit complex [36], the kinetoplastid *Trypanosoma cruzi* has a 12-subunit complex [37], and in the ciliate *Tetrahymena* this complex consists of 15 subunits [38]. Recent studies in apicomplexans found an 8-9 subunit complex in *Plasmodium* and *Toxoplasma,* respectively [10–13], and its enlargement in *Eimeria* was also suggested [16]. All complex II compositions possess two conserved matrix-facing subunits, SDHA and SDHB, that form the soluble enzymatic domain that contains the FAD and Fe-S cofactors, as well as between 2 and 13 membrane-embedded subunits that anchor the complex to the IMM. In the “canonical” enzyme those subunits are named SDHC and SDHD and they further house the ubiquinone binding site. The amino acids involved in ubiquinone binding in eukaryotes with enlarged subunit number have not yet been experimentally characterized.

Searching our mitochondrially-enriched complexome dataset for the profiles of homologs of the highly conserved SDHA and SDHB subunits identified proteins with a peak at both high molecular weight (∼900-990 kDa) and low molecular weight (between ∼60 and 120 kDa) (Fig 2A, S4, S5 Tables). The low-molecular-weight peak likely reflects either an assembly intermediate of the soluble matrix domain or subunit monomer migration, while the one occurring at high molecular weight represents the mature complex. This size of complex II is consistent with the BN-PAGE in-gel stain (Fig 1B). The whole-cell complexome dataset also revealed evidence of a sub-assembly containing SDHA and SDHB migrating at ∼320-350 kDa (Fig 2B). Within the enriched mitochondrial dataset, 6 proteins that are homologs of the recently identified apicomplexan membrane-anchoring complex II subunits (Fig 2A) peak at ∼900 kDa. In the whole-cell complexome, these subunits migrated at high molecular weight of ∼1290 kDa (Fig 2B). These findings provide support for an affiliation of these 6 proteins as subunits of complex II in *Perkinsus*. Their assignment to complex II is further supported by a recent structural study of the mETC supercomplex in *Perkinsus* that identified them as part of a 10-subunit complex II [39].

**Figure 2:**
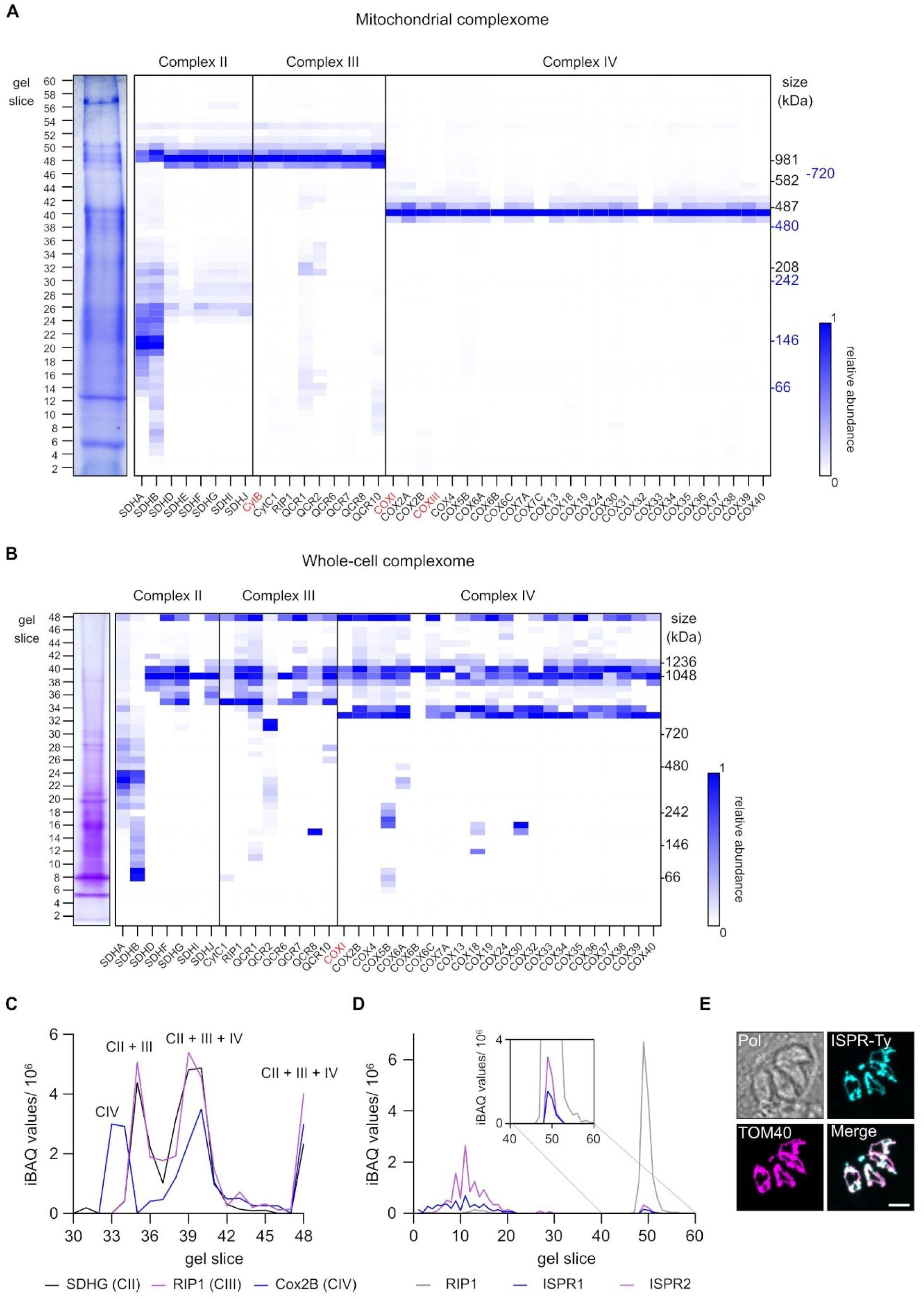
Complexome profiles of mETC complexes. (A) Heatmaps of the complexome profiles from mitochondrially-enriched material of mETC complex subunits from complex II, complex III and complex IV. Each row represents a gel slice, with the gel slice number indicated on the left and the molecular weight, based on the migration of mammalian mitochondrial complexes (black) and soluble proteins (blue), on the right. Subunit IDs are shown at the bottom of their corresponding profile, with mitochondrially-encoded subunits highlighted in red. Dark blue indicates the highest relative abundance (1) and white the lowest (0). Coomassie-stained Native PAGE gel on which the analysis was performed is depicted on the left of the profile. Data shown is from replicate 1. (B) Heatmap of the whole-cell complexome of mETC complexes, as in part *(A)*. Molecular weight, based on the migration of soluble proteins on the right is shown on the right. (C) Abundance profile of complex II subunit SDHG, complex III subunit RIP1 and complex IV subunit Cox2B across all gel slices. Protein abundance is displayed as iBAQ values. Data shown is from whole-cell complexome. (D) Abundance profile of ISPR1, ISPR2 and complex III subunit RIP1 across all slices. Protein abundance is displayed as iBAQ values. Data shown is from replicate 2. (E) Immunofluorescence assay of *Toxoplasma gondii* transiently expressing a Ty-epitope tagged version of *Tg*ISPR. Samples labelled with anti-Ty to detect *Tg*ISPR-Ty (cyan) and anti-Tom40 (magenta) to detect the mitochondrial marker protein Tom40. Scale bar is 5 µM.

We next searched for complex III subunits. In the mitochondrially-enriched complexome dataset, the mitochondrially-encoded cytochrome *b* subunit was found with a peak at ∼900 kDa (Fig 2A). Among other proteins with a peak in the same slice, we found 8 other putative complex III subunits, identified based on the homology to known apicomplexan subunits and to those identified in the recent mETC supercomplex structure [39]. Searching the whole-cell complexome dataset identified 8 of these subunits as migrating in the same gel slice providing additional support to their affiliation (Fig 2B). Notably, in both datasets we found subunits of both complexes II and III predominantly migrating in the same gel slice.

Additionally, we observed Pmar_PMAR021073, a predicted homolog of the complex III assembly factor BCS1, with a peak in the gel slice below mature complex III, at ∼820 kDa, and with high levels also within the complex III gel slice (S1 Table). In other systems, BCS1 is involved in the maturation of the Rieske (RIP1) subunit of complex III and is found in a late stage of assembly [40], suggesting that in our mitochondrially-enriched complexome, we observe a late assembly intermediate as well as mature complex III. Evidence of a lower molecular weight sub-assembly of complex III subunits QCR1 and QCR2 are seen in both datasets as well (Fig 2A,B).

Complex IV has shown considerable variation in subunit number across the eukaryotes. Apicomplexans possess an enlarged complex with 25 subunits, compared to just 14 observed in mammals [9]. The recent *Perkinsus* mETC supercomplex structure suggests a 27-subunit complex [39]. We first searched for the mitochondrially-encoded CoxI and CoxIII subunits and found them with a peak in a gel slice representing ∼420 kDa (Fig 2A; S4 S5 Tables). Moreover, this size of complex IV is consistent with the BN-PAGE in-gel stain (Fig 1B). We detected 24 of the expected subunits in the same gel slice, thereby confirming complex IV’s composition (Fig 2A). Interestingly, we were not able to detect Cox2c, which was observed in the *Perkinsus* structure, and its homolog was found in a recent structural study from *Toxoplasma* [9]. However, this subunit has not been detected in numerous proteomic studies of the apicomplexan mETC [3,15,28], suggesting challenges in its detection by mass spectrometry. Searching the whole-cell complexome dataset identified CoxI and 21 further complex IV subunits, migrating at high molecular weights and with up to three peaks of abundance, likely relating to multimers or supercomplex formation (see below) (Fig 2B).

Associations between individual mETC complexes, termed supercomplexes, have been observed in numerous systems, recently including Apicomplexa [9]. Likewise, the above-mentioned *Perkinsus* structural study described a supercomplex composed of complexes II, III and IV (II2-III2-IV2) [39]. We interrogated our datasets to see if we could detect any supercomplex. In the complexome from mitochondrially-enriched samples, complexes II and III are found co-migrating, but subunits of complex IV are not present in this gel slice. This is not unexpected, as the βDDM extraction does not typically retain weaker inter-complex associations. The association of complexes II and III could be due to the tight binding conferred by a long helical extension in SDHG observed in the structure [39]. We therefore interrogated our whole-cell dataset, extracted in 0.1 % IGEPAL, which is more likely to retain weaker inter-complex interactions. Here we observed that subunits from complexes II, III and IV indeed appear together in multiple peaks of high relative abundance (Fig 2B, C). For example, complex IV subunits have peaks at ∼760-840, ∼1,300-1,400 and ∼2,800 kDa. Moreover, the peaks from different respiratory complexes overlap, consistent with the detection of a supercomplex containing complexes III and IV (Fig 2C).

Our complexome datasets allow identification of almost complete mETC complexes, validating their composition, and demonstrating that the enlarged mETC complexes observed in Apicomplexa are conserved across a broader group within the myzozoans and thus likely evolved in their common ancestor.

### Evidence of interaction between complex III and putative inhibitors

A structural study of the *Perkinsus* mETC supercomplex [39] identified two novel proteins that interact with the RIP1 subunit of complex III. These proteins were proposed to act as inhibitors of the complex’s enzymatic activity due to their interaction site overlapping with the binding site for cytochrome *c*. These proteins were named ISPR1 and ISPR2 (for Iron-Sulfur Protein Related). To lend support to this observation we searched for them in our complexome datasets. Both proteins were detected in both replicates of our complexome on mitochondrially-enriched samples. Major peaks were identified at low molecular weight slices likely representing free monomers or dimers. However, in the second replicate a secondary, smaller, peak was detected in gel slices that contains complex III subunits (Fig 2D), consistent with an interaction between ISPR1, ISPR2 and this complex.

The identification of a novel complex III inhibitory factor in *Perkinsus* raises the question of whether it is a lineage-specific innovation, or if it is more widely conserved. It was noted that *Toxoplasma* contains a likely homolog, TGGT1_224932 [39]. To test if the mitochondrial localisation of this putative *Tg*ISPR is conserved, we exogenously expressed a Ty epitope-tagged version of the protein in *Toxoplasma* under the tubulin promoter and performed immunofluorescence analysis. We observed signal with a characteristic mitochondrial “lasso-shape” morphology [41], which co-localised with the mitochondrial marker TOM40 [42], supporting mitochondrial localisation (Fig 2E). Future functional studies will be required to determine whether *Tg*ISPR performs an inhibitory role as proposed for *Perkinsus*. Overall, our results lend support to the observation that novel proteins interact with complex III in *Perkinsus* and lays the foundation for future functional investigations in the apicomplexan models.

### Detection of F_1_F_o_-ATP synthase

In *Toxoplasma* F_1_F_o_-ATP synthase complex is a huge 1.85 MDa complex consisting of 32 subunits [10,17–19], compared to ∼15 canonical subunits of the yeast and mammalian systems. Immunoblot analysis (Fig 1B) provides hints that *Perkinsus* possesses a similarly enlarged ATP synthase. Searching for homologs of confirmed *Toxoplasma* ATP synthase subunits within the *Perkinsus* genome identified 27 putative homologs (S4 Table). Of these, 26 were present in our mitochondrially-enriched complexome dataset and 25 in the whole-cell dataset. The conserved subunit c is an expected exception, as it has previously proved recalcitrant to detection by mass spectrometry [18]. All 26 subunits peaked in relative abundance at ∼1.9 MDa in the mitochondrial complexome (Fig 3A; S4, S5 Tables), and at ∼1.5 MDa in the whole-cell complexome (Fig 3B), confirming an enlarged ATP synthase. Conserved subunits include ATPTG8 and 9, which contain coiled-coil-helix-coiled-coil-helix (CHCH) domains. Subunits containing CHCH-domains have recently been shown to be essential for ATP synthase formation [43] and were previously only studied in apicomplexans. Their presence in *Perkinsus* suggests CHCH-domain containing ATP synthase subunits are a myzozoan-wide feature. The ATP synthase inhibitor subunit IF1 was found also in a secondary low-molecular-weight peak, consistent in size with monomer or dimer migration, and providing evidence that the IF1 pool is not permanently engaged with the mature complex. Likewise, the conserved catalytic subunit ATPβ also has a low molecular weight peak, matching the secondary band seen by immunoblot analysis (Fig 1B).

**Figure 3:**
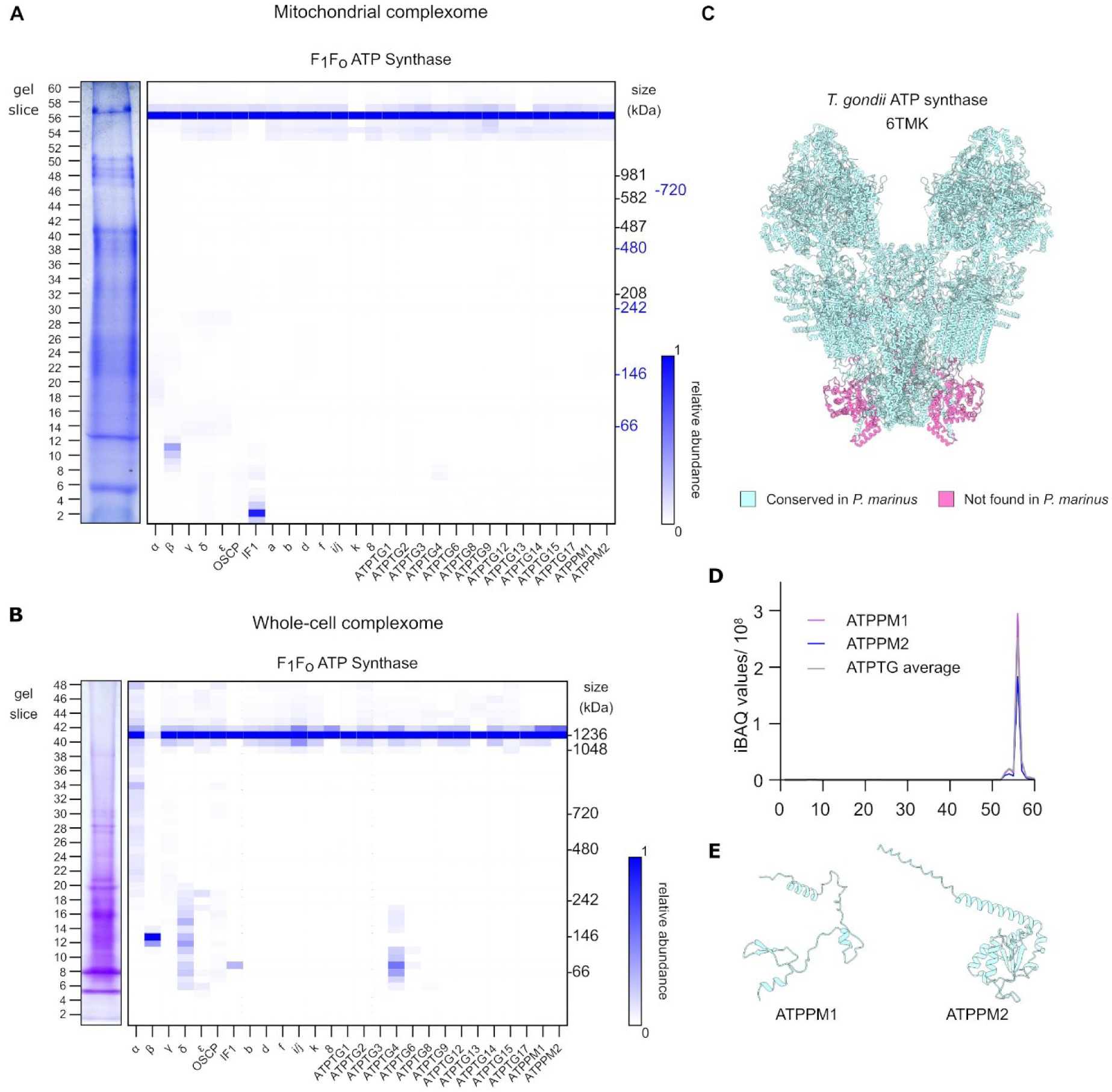
Complexome profiles of the F_1_F_o_ ATP synthase complex. (A) Heatmap of the complexome profile from mitochondrially-enriched material of ATP synthase. Each row represents a gel slice, with the gel slice number indicated on the left and the molecular weight, based on the migration of mammalian mitochondrial complexes (black) and soluble proteins (blue), on the right. Subunit IDs are shown at the bottom of their corresponding profile. Dark blue indicates the highest relative abundance (1) and white the lowest (0). Coomassie-stained Native PAGE gel on which the analysis was performed is depicted on the left of the profile. Data shown is from replicate 1. (B) Heatmap of the whole-cell complexome of ATP synthase, as in part *(A)*. Molecular weight, based on the migration of soluble proteins on the right is shown on the right. (C) Structure of ATP Synthase dimer from *Toxoplasma gondii* (PDB 6TMK, [17]) with subunits conserved in *Perkinsus* coloured blue, and subunit not found in coloured pink. Visualised using ChimeraX [86]. (D) Abundance profile of ATPPM1, ATPPM2, and the average abundance of all ATPTG subunits across all gel slices. Protein abundance is displayed as iBAQ values. Data shown is from replicate 1. (E) Structural prediction of ATPPM1 and ATPPM2 (predicted using Alphafold3 [87]).

We were unable to identify *Perkinsus* homologs of many of the lineage-specific lumen-facing subunits identified from the *Toxoplasma* structure (ATPTG5,7,10,11,16) (Fig 3C). *Perkinsus* may lack an expanded luminal domain, or possesses derived versions of the missing subunits, which we were not able to detect by homology searches due to sequence divergence. To explore further we searched our complexome for co-migrating proteins which may be novel subunits. We noted that, in all complexome datasets, 2 other proteins (Pmar_PMAR021141 and Pmar_PMAR014495) consistently co-migrated with the other subunits of ATP synthase, with similar abundance to other myzozoan-specific ATP synthase subunits (Fig 3A, B, D). Similar to the missing subunits from *Toxoplasma* ATPTG10,11 and 16, Pmar_PMAR021141 is a short protein of 10 kDa. Pmar_PMAR014495 is a 19 kDa protein with a predicted thioredoxin domain (Fig 3E). To investigate further, we first purified ATP synthase by size exclusion chromatography and submitted it for mass spectrometry analysis (S2A, B Fig). In addition to detection of 22 expected ATP synthase subunits, both of these new proteins were also present (S6 Table). These data suggest that they are genuine ATP synthase subunits, and we putatively assign them the names ATPPM1-2, although further experimental work would be required to confirm this. Combined, these data confirm that *Perkinsus* has an enlarged ATP synthase, and that this divergent complex is conserved across sampled myzozoans and reveals putative new species-specific subunits.

It was previously observed in *Toxoplasma* that the gene encoding ATP synthase subunit ATPTG1 also separately encodes the complex III subunit CytC1, suggesting post-translational cleavage [10] and potentially coupled assembly of the mETC and ATP synthase [44]. Homology analysis showed that this fusion was conserved across a subset of apicomplexans and that outside this group ATPTG1 and CytC1 are encoded by separate genes [10] (S3C, D, S4 Figs). Our detection of ATPTG1 in complex with ATP synthase strongly suggests a role for this protein as an ATP synthase subunit prior to the fusion event with CytC1.

### Mitochondrial ribosome

The mitoribosome is a macromolecular complex required to translate transcripts encoded by the mitochondrial genome, consisting of two large riboprotein subunits – the large (LSU) and small subunit (SSU). Both the LSU and SSU are composed of numerous proteins and rRNAs. In the myzozoan lineage, the mitochondrial genomes predict an unusually fragmented rRNA backbone [5,20,21,45], which would require the function of new proteins to support translation. Two recent structural studies of *Toxoplasma* confirmed this hypothesis and demonstrated that, similarly to mETC and ATP synthase, apicomplexans have mitoribosomes that diverge significantly from their better studied homologs in humans and yeast [23,24]. Notably, the *Toxoplasma* mitoribosome contains at least 124 subunits (73 in LSU, 51 in SSU), including 55 clade-specific ones [23]. In contrast, the yeast and human mitoribosomes contain in total only 75 and 80 subunits, respectively [46,47]. This difference supports the notion that the recruitment of this group of unorthodox mitoribosomal proteins acts to mitigate the highly fragmented rRNA.

To test if this divergent composition is conserved across myzozoans, we first searched for homologs of confirmed *Toxoplasma* mitoribosomal proteins within the *Perkinsus* genome, and identified 78 putative homologs (S4 table). Of these, 74 were identified in at least one complexome dataset. In the first replicate of the mitochondrial complexome, 70 subunits were detected, with 67 of them showing their highest abundance in the top gel slices (∼ 2 MDa) (Fig 4). The LSU and SSU subunits were observed most prominently in the top two and three gel slices, respectively. This slight difference in migration could represent separate migration of the LSU and SSU, or a mixture between full monosomes and individual subunits. We observed 21 of the 55 clade-specific subunits co-migrating with the mitoribosome, suggesting partial conservation across myzozoans. In the whole-cell complexome, we detected 18 out of 74 putative subunits, 5 from the SSU and 13 from the LSU (S5 Table). The lower number of detected proteins potentially reflects different detergent extraction conditions used in the mitochondrial dataset, leading to the differential preservation of higher-order molecular interactions. As observed in the mitochondrial dataset, the LSU and SSU proteins possess slightly different migration profiles, with 7 out of 14 LSU proteins showing their highest peak at ∼1.7 MDa, while 4 out of 5 detected SSU factors migrate mainly above 2 MDa.

**Figure 4:**
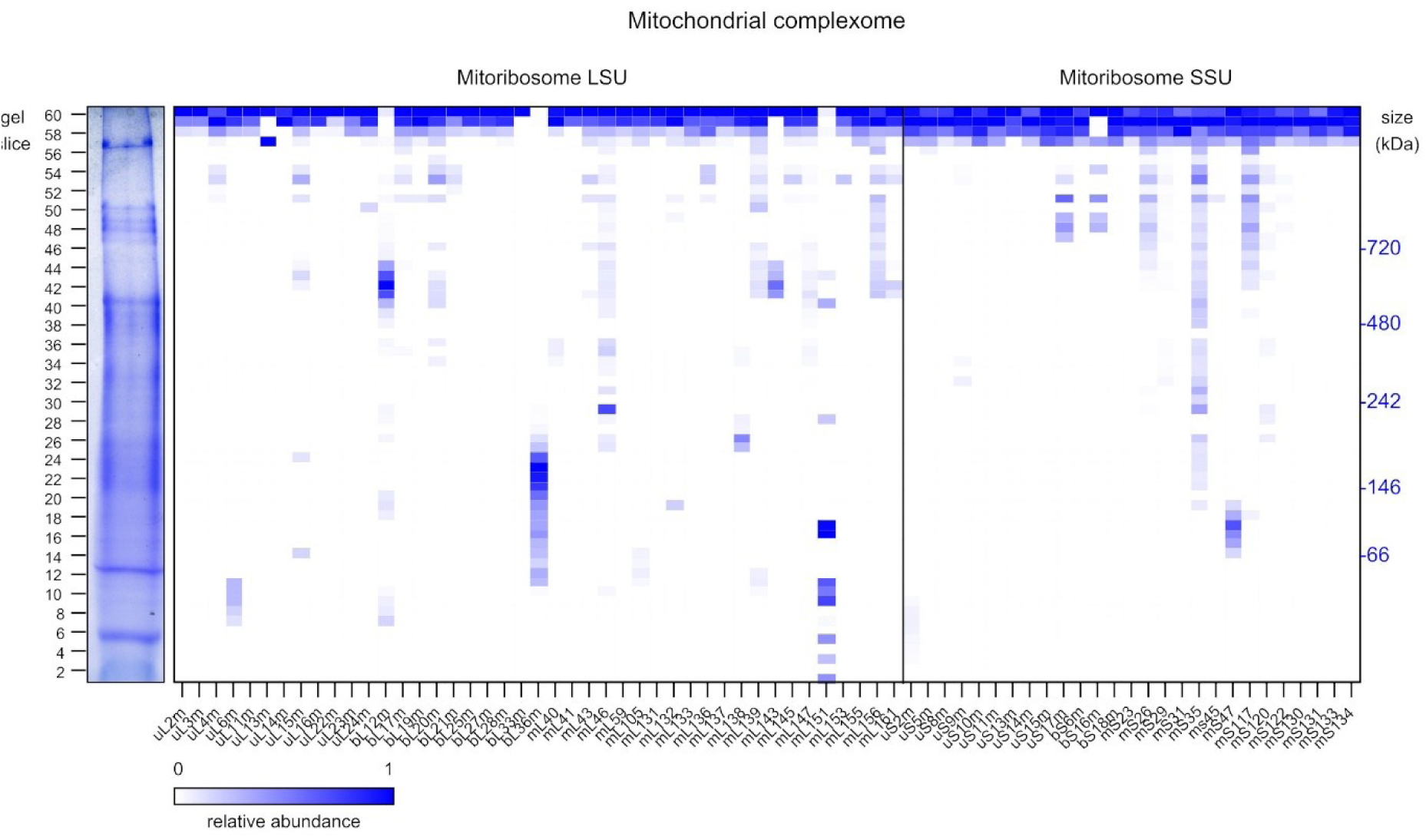
Complexome profile of mitoribosome. Heatmap of the complexome profile of the mitochondrial ribosome, showing subunits of the LSU and SSU. Each row represents a gel slice, with the gel slice number indicated on the left and the molecular weight, based on the migration of mammalian mitochondrial complexes (black) and soluble proteins (blue), on the right. Subunit IDs are shown at the bottom of their corresponding profile. Dark blue indicates the highest relative abundance (1) and white the lowest (0). Coomassie-stained Native PAGE gel on which the analysis was performed is depicted on the left of the profile. Data shown is from replicate 1.

Structural analyses of the *Toxoplasma* mitoribosome revealed the extent of rRNA fragmentation and provided insight into the mechanism through which lineage-specific proteins mitigate this fragmentation and promote ribosomal function through RNA-protein interactions [23]. For example, 11 *Toxoplasma* proteins interact each with multiple rRNAs and therefore were assigned a putative role as “joints”. *Perkinsus* is also thought to contain highly fragmented rRNAs [25]. We detected 8 of the 11 joint proteins, suggesting conservation in the architecture of protein scaffolds for supporting rRNA fragment stability and formation of critical mitoribosomal functional domains. This included a RAP (**<u>R</u>**NA-binding domain abundant in **<u>ap</u>**icomplexans) domain-containing protein homologous to *Toxoplasma* mL138, which is essential for mitoribosome stability [23]. Five RAP domain proteins were found in the *Toxoplasma* mitoribosome with this suggested regulatory role, three of which (mL136, mL138 and mL147) are conserved in *Perkinsus* and migrate with the mitoribosome.

Myzozoan genomes encode proteins with the AP2 transcription factor domains. In the apicomplexans, numerous members of this family (called ApiAP2) are involved in controlling gene expression-dependent life cycle stage transitions [48]. Surprisingly, in *Toxoplasma,* 4 out of 67 ApiAP2 proteins, have been incorporated into the mitoribosome [23,24]. *Perkinsus* has 25 predicted ApiAP2 proteins[49], 4 of which migrate with the mitoribosome (mS130, mL143, mL153, mL161) (Fig 4; S5 Table). This suggests that the unusual incorporation of AP2-like proteins in the mitoribosome is conserved across the myzozoans. Additionally, the *Toxoplasma* mitoribosome contains proteins with domains that may indicate repurposing from another function. Two examples include mL151, which has a BolA-like domain, commonly found in the iron-sulfur cluster assembly pathway proteins, and mS134, which has an acylphosphatase enzymatic domain. Both proteins appear conserved as mitoribosomal components in *Perkinsus* as their homologs peak together with the other mitoribosomal proteins.

Overall, several unusual features recently observed in *Toxoplasma* mitoribosome appear conserved in *Perkinsus*, suggesting that these innovations are Myzozoa-conserved. Future structural studies will be needed to unravel the full extent of conservation of these features, and any unique attributes of the *Perkinsus* mitoribosome.

### Complexes from other cell compartments

Finally, we explored whether our complexome datasets could provide insight into other cell compartments. We reasoned that our whole-cell complexome would provide cell-wide coverage, and, as previously highlighted, immunoblot analysis of fractions from our mitochondrial purification protocol suggested co-purification of other organelles (Fig 1A). We first investigated the 26S proteasome complex. Homology searches identified *Perkinsus* homologs of 32 out of 33 subunits from the human complex [50]. We could identify 17 subunits of the 19S proteasome/ regulatory particle migrating at 1.4 MDa in our mitochondrial complexome, and 16 subunits migrating at 1.2 MDa in our whole-cell complexome (Fig 5A, B). Similarly, we could detect 14 subunits of the 20S proteasome/core particle migrating at 690 kDa in our mitochondrial complexome and 13 subunits migrating at 765 kDa in our whole-cell complexome. Detection of the proteasome was also observed in a previous mitochondrial complexome of *Plasmodium falciparum* [11].

**Figure 5:**
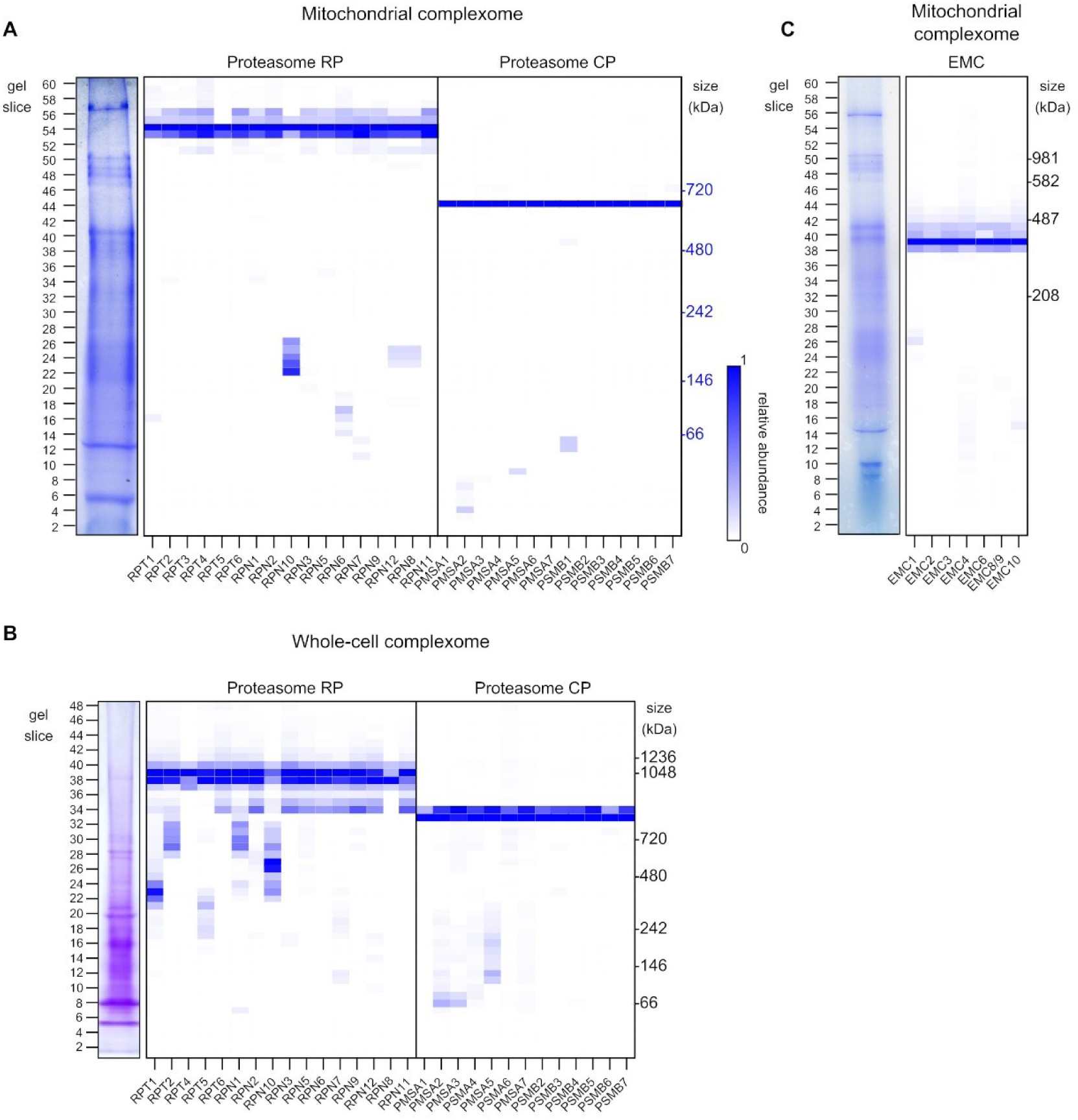
Complexome profile of non-mitochondrial complexes. (A) Heatmap of the complexome profile from mitochondrially-enriched material of the 26S proteasome, comprised of the 19S regulatory particle (RP) and the 20S core particle (CP). Each row represents a gel slice, with the gel slice number indicated on the left and the molecular weight, based on the migration of soluble proteins (blue), on the right. Subunit IDs are shown at the bottom of their corresponding profile. Dark blue indicates the highest relative abundance (1) and white the lowest (0). Coomassie-stained Native PAGE gel on which the analysis was performed is depicted on the left of the profile. Data shown is from replicate 1. (B) Heatmap of the whole-cell complexome of the 26S proteasome, comprised of the 19S regulatory particle (RP) and the 20S core particle (CP), as in part *(A)*. Molecular weight, based on the migration of soluble proteins on the right is shown on the right. (C) Heatmap of the complexome profile from mitochondrially-enriched material of the Endoplasmic reticulum membrane protein complex (EMC), as in part *(A)*. Molecular weight is based on the migration of mammalian mitochondrial complexes. Data shown is from replicate 2.

Since immunoblot analysis suggested co-purification of the endoplasmic reticulum, we searched our mitochondrial complexome for the endoplasmic reticulum membrane protein complex (EMC). We identified in *Perkinsus* clear homologs of 7 of the 9 human subunits [51], which migrated at 330 kDa in our mitochondrial complexome (Fig 5C). Recently, this complex was also identified in the *Plasmodium* complexome [11]. Interestingly, unlike in *Plasmodium*, we were able to detect the EMC6 subunit migrating with other EMC components.

Detection of the proteasome and EMC suggests our datasets are suitable for exploring the composition of macro-molecular protein complex in other cellular compartments, and may provide extra insight and comparison to future spatial proteomic studies.

## Discussion

Mitochondria are unique and defining features of the eukaryotic cell and play a variety of important roles in their function. However, our knowledge of how they operate is largely derived from a narrow set of model organisms that are often closely related. Indeed, the vast majority of eukaryotic diversity, especially in mitochondrial biology, is “hidden” in unicellular eukaryotes and, despite increased recent research efforts, we still know surprisingly little about their molecular features.

An example of the diversity of mitochondrial functions across the eukaryotes is in the variety of mitochondrial proteomes. The size of an organism’s mitochondrial proteomes is often indicative of the organism’s mitochondrial capacity [52]. Unicellular eukaryotes offer examples from both ends of this spectrum. For example, the most complex mitochondrial proteomes to date come from the euglenid *Euglena gracilis* and *Paradiplonema* (alternatively referred to as *Diplonema*) *papillatum* with estimated ∼1,800 and ∼1,900 proteins [53]. At the other end of the spectrum, the mitochondria of other protists have been subject to massive reduction [54], including some that have dispensed with mitochondria altogether [55]. Indeed, parasitic lineages are well known for streamlining their genomes [56], which likely provides spatial efficiency and energy conservation [57], with the mitochondria being a clear example of this trend. The greatest diversity of mitochondrial biology is therefore found in unicellular eukaryotes [58].

One area in which this diversity manifests itself is in the composition and features of the mitochondrial complexes involved in oxidative phosphorylation. Over the past few years, a number of studies in a variety of protist lineages have shown considerable divergence in the composition and features of these mitochondrial complexes in comparison to the “textbook” model systems, such as yeast and mammals. A stark example comes from recent structural studies of the ciliate *Tetrahymena thermophila* that demonstrated a respiratory chain composed of ∼150 proteins compared to ∼90 in humans [38,59–61]. Similar findings have been found in the apicomplexans. The mitochondrial complexes of this lineage have been the subject of sustained research effort in the past decade, largely due to their global importance as pathogens of humans and livestock, and their development as powerful tractable model organisms. Through both proteomic and structural approaches, it has been shown that almost every mitochondrial complex is substantially larger than the opisthokont version, contrary to the simplified yet prevailing notion that parasite pathways are reduced compared to their host. These observations span complex II [10–13], complex IV [9–11,15]and ATP synthase [10,11,17–19]. The larger size is an outcome of numerous species or clade-specific subunits, and the field is still exploring the driving forces for their recruitment.

Despite this progress, it was still unclear if the newly discovered subunits were specific to the apicomplexans, or more widely conserved across other related lineages within the myzozoa. Here, we aimed to answer this question by exploring the mitochondrial complexes of an early-diverging myzozoan, *P. marinus*. Our focus on the Perkinsozoa clade, which is sister group to the dinoflagellates, and branched from the lineage that forms apicomplexans and chromerids early in myzozoan evolution, allows insight into the evolution of these features from the ancestral state. Shared mitochondrial features found here would suggest their emergence prior to the split and, importantly, prior to the shift to parasitism in Apicomplexa.

Our findings broadly point to a conservation of mitochondrial complex composition both for the oxidative phosphorylation complexes and the mitoribosome. Novel innovations first observed in the apicomplexan parasites, for example the enlarged subunit composition of complexes II and IV, and ATP synthase, as well as the addition of novel proteins to the mitoribosome, are observed in *Perkinsus*, consistent with conservation across myzozoans. This suggests a scenario where the last common ancestor of the 4 diverging lineages, namely dinoflagellates, chromerids, apicomplexans and perkinsozoans, already possessed enlarged mitochondrial complexes consisting of many of the clade-conserved and -specific subunits seen in both parasites. Given the number of free-living species in these lineages, and the multiple independent origins of parasitism, this would also imply this feature is unrelated to a parasitic lifestyle.

A question that naturally arises is how these enlarged complexes in unicellular eukaryotes arose and what purpose the additional subunits play. We do not have definitive answers, but several reasons can be discussed. In some cases, numerous subunits with reduced hydrophobicity provide the structural features of a single more hydrophobic subunit, and it has been hypothesised that reduced mitochondrial genome content is a driving force for this outcome [9,17,39,62], as they allow transfer of subunit-encoding genes from the mitochondrial to nuclear genome [63]. This process may be aided by splitting up of large hydrophobic proteins, that would prove problematic for mitochondrial import, into more numerous, less hydrophobic, subunits [10]. Another theory is that these extra subunits imbue the complex with additional functions. Structural studies revealed that, in many cases, species-or clade-specific subunits mediate interactions between complexes and multimerization of complexes, which seem to evolve in line with the shape of mitochondrial cristae [9,17,38]. An example of this is the additional subunits in *Toxoplasma* ATP synthase, which allow the formation of hexamers, with a role in shaping the cristae membrane [17]. Likewise, in the formation of *Toxoplasma* supercomplex, complexes III and IV interact in an alternate orientation from mammals, and this novel interaction interface is lined with the lineage-specific subunits [9]. In *Perkinsus*, the novel subunit, SDHG, appears to promote close associations between complexes II and III, perhaps aiding in supercomplex stability [39]. Extra subunits have also been proposed to play a role in maintaining complex stability in mitoribosomes [23,64,65]. The reason for a largely increased repertoire of accessory subunits may be a combination of (all) the above.

Our study revealed a number of components conserved between the *Perkinsus* and the *Toxoplasma* mitoribosomes, indicating that they potentially share many of the unusual features recently uncovered in the apicomplexans [23,24]. The unique mitoribosomal rRNA fragmentation is common to all myzozoans whose mitochondrial genomes have been sequenced [22,25,66] and, therefore, it is likely that features that evolved to “mend” the fragments, control assembly, and maintain stability are also shared. Structural observations derived from the *Toxoplasma* mitoribosome, the only one available for myzozoans, pointed out three main features that have the potential to mitigate the extreme rRNA fragmentations: i) proteins that binds several rRNA molecules and thus act as “joints”; ii) high abundance of short peptides that are enriched with aromatic residues capable of stacking with nucleobases and with charged residues able to interact with backbone rRNA, thus promoting association between rRNA molecules; and, iii) the addition of poly-A tails onto rRNA that are bound by proteins, which in return bind each other, promoting indirect association between rRNAs. The *Perkinsus* mitoribosome peak contains homologs of “joint” proteins, but we were unable to identify homologs of the short peptides which promote association of short rRNAs. This may imply a different mitigation solution for dealing with short rRNAs has evolved in *Perkinsus*. Alternatively, short peptides may be present that have extensively diverged from *Toxoplasma* making them hard to detect by homology searches. Likewise, homologs of the proteins that bind to poly-A tails migrate with other *Perkinsus* mitoribosomal proteins suggesting that this mechanism is also conserved. This is further supported by the identification of a homolog of TgmtPAP1, which is a mitochondrial poly-A polymerase important for the complete polyadenylation of mitoribosomal rRNA in *Toxoplasma* [23].

The observation that 4 members of the ApiAP2 transcription factor family are integral mitoribosome components in *Toxoplasma* [23,24] is extended here to *Perkinsus*. However, while *Toxoplasma* has an unusually expanded number of apiAP2 family members, 67 in total, the *Perkinsus* genome encodes only 25 [49], and 4 of them are mitoribosomal. Homologs of those 4 ApiAP2 are found across the myzozoan clade including dinoflagellates, in which 5 have been detected. It is tempting to hypothesise that rRNA fragmentation and the acquisition of these 4 mitoribosomal proteins with ApiAP2 folds have co-evolved; however, this requires further investigation.

Recent studies on the *Perkinsus* mitochondrial genome highlighted its several specific features, including replacement of the canonical UGA stop codon to code for tryptophan, and an unusual programmed ribosomal frameshifting found in all genes, which seems to require 18 to 19 corrections across the encoded mitochondrial proteins [25]. These irregularities would necessitate adjustment in mitoribosome functional control, and potentially result in the evolution and recruitment of *Perkinsus*-specific proteins and features. We found numerous proteins, annotated as uncharacterised, co-migrating with confirmed mitoribosomal proteins. Although we cannot currently provide support for their mitoribosome affiliation, as some may belong to other high molecular complexes, it is likely that many are *bona fide Perkinsus*-specific mitoribosome proteins. Structural studies would assist in assigning any new proteins and revealing any *Perkinsus*-specific mitoribosome features.

*Perkinsus marinus* has proved a versatile model organism for understanding myzozoan biology and is increasingly being utilised for investigations into divergent eukaryotic biology, of which this study is an example. We applied a proteomic complexome profiling approach [27], which has proved a powerful method for uncovering the composition of organellar complexes [28,29]. It has most frequently been used for studying organelles in more commonly studied organisms, and significantly contributed to our understanding of pathologies in mitochondrial disease patients [67], mammalian [68] and yeast mitochondrial biology [69], and plant and algal organellar biology [70,71]. More recently, it has also been successfully used for exploring non-model lineages. Recent complexome studies of unicellular eukaryotes include *Plasmodium* [11], *Toxoplasma* [10], the extremophile yeast *Rhodotorula mucilaginosa* [72] and this study. We believe complexome profiling will be a powerful tool for future explorations of divergent mitochondrial biology.

## Materials and methods

### Cell culture

*Perkinsus marinus* cells (strain ATCC 50983), used the mitochondrially-enriched complexome, were grown in liquid culture in ATCC medium 1886 supplemented with penicillin and streptomycin. *Perkinsus marinus* strain PRA-240 (CB5D4; [31]), used for the whole-cell complexome, was grown in modified DME:Ham’s F12 (1:2) medium supplemented with 5 % foetal bovine serum, 50 mM HEPES, 3.6 mM sodium bicarbonate, 15 g/L artificial sea salts, 100 U/mL penicillin G, and 100 U/mL streptomycin sulphate, pH 6.6. Cells were grown without agitation in the dark at 25-26 °C. For culture maintenance, 200 µL of stationary-phase culture were transferred to a T25 culture flask with 10 mL of fresh medium once a week.

### Isolation of mitochondrial material

Enriched mitochondrial fractions were obtained by cell lysis followed by differential centrifugation. *Perkinsus* cells were grown in ∼1.5 L of liquid culture and harvested at OD_600_ ∼1. Cells were pelleted by centrifugation at 1000 x g for 15 minutes at 4 °C and weighed. Per gram of cells, 2 g fine glass beads and 2 mL ice-cold mito-membrane buffer (0.6 M sucrose, 20 mM MOPS-NaOH, pH 7.5, 1 mM EDTA) containing protease inhibitors were added. Cells were disrupted by 8 rounds of 1-minute vortexing and 1-minute incubation on ice. Differential centrifugation was performed at 3,500 × *g* for 10 minutes at 4 °C followed by 40,000 × *g* for 120 minutes at 4 °C. Pellets containing mitochondrial membranes were stored at -80 °C before use.

### Preparation of whole-cell samples

5 x 10^7^ of cells (OD_600_ ∼2) were harvested by centrifugation at 1,700 × *g* for 10 min at 4°C. The cell pellet was resuspended in 50 µl of solubilization buffer (50 mM NaCl, 50 mM Bis-Tris/HCl pH 7.0, 2 mM 6-Aminocaproic acid (ACA), 1 mM EDTA) containing 0.1% IGEPAL CA-630 detergent and supplemented with 1x complete EDTA-free protease inhibitor and 12.5 µg of 0.1% (w/v) DNase. 1 g of fine glass beads were added, and cells were disrupted by vortexing for 3 min at 4°C. To remove unlysed cells and insoluble material, the cell lysate was cleared by centrifugation at 1,000 × *g* for 10 min at 4°C. Protein concentration was determined by the BCA protein quantification assay.

### Blue native, SDS PAGE and immunoblot analysis

#### SDS-PAGE

7.5 µg of protein lysate, measured using a Bradford assay, was resuspended in Laemmli buffer (2% (w/v) SDS, 125 mM Tris-HCl, pH 6.8, 10% (w/v) glycerol, 0.04% (v/v) β-mercaptoethanol, 0.002% (w/v) bromophenol blue) and heated at 95 °C for 5 min. Proteins were then separated on a 12.5 % SDS-polyacrylamide gel. EZ-Run Prestained Rec protein ladder was used as a molecular weight marker. Proteins were transferred under semi-dry conditions to a nitrocellulose membrane and labelled with primary antibodies: anti-ATPβ (Agrisera AS05085, 1:5000), anti-histone H3 (Abcam ab1791, 1:1,000), anti-GAPDH (Abcam ab125247, 1:2,000), anti-BiP (a gift from James D. Bangs, 1:5,000). This was followed by labelling with secondary fluorescent antibodies IRDye 800CW, 680RD (1:10,000, LIC-COR) and detection with an Odyssey CLx. As a loading control, duplicate gels run in parallel were incubated for 1 hr in instant blue.

#### Blue Native (BN) PAGE for mitochondrially-enriched samples

Samples were resuspended in solubilization buffer containing 1.75 % βDDM, 750 mM aminocaproic acid, 50 mM Bis-Tris-HCl (pH 7.0), 0.5 mM EDTA. This was followed by incubation on ice for 30 min followed by centrifugation at 16,000 × *g* at 4°C for 30 min to separate solubilized protein complexes from insoluble material. This supernatant was combined with sample buffer containing Coomassie G250, resulting in a final concentration of 0.4375% βDDM and 0.1% Coomassie G250 and a protein: detergent ratio of ∼1: 5.8. 15-30 µg of sample was separated on a 4-16% Bis-Tris gel under native conditions. Bovine mitochondrial membranes, as well as NativeMark were used as a molecular weight marker.

To visualize total proteins, gels were incubated in a 0.1% (w/v) Coomassie R-250, 45% methanol (v/v), 10% acetic acid (v/v) solution, followed by de-staining and incubation in milliQ water. For immunoblot analysis protein was transferred to a PVDF membrane and probed with anti-ATPβ as above. In-gel histochemical stains were performed as in [73]. Briefly, for detection of succinate dehydrogenase the gel was incubated in 50 mM KH_2_PO_4_, pH 7.4, 0.1 mM ATP, 0.2 mM phenazine methosulfate, 10 mM succinate and 0.2% (w/v) nitroblue tetrazolium until the stain was observed, and the reaction stopped and gel further destained by incubation in 45% methanol (v/v), 10% acetic acid (v/v). For detection of cytochrome *c* oxidase activity, the gel was incubated with 50 mM KH_2_PO_4_, pH 7.2, 1 mg ml−1 cytochrome *c*, 0.1% (w/v) 3,3′-diaminobenzidine tetrahydrochloride until the stain was observed, and the reaction stopped and gel further destained by incubation in 45% methanol (v/v), 10% acetic acid (v/v).

For complexome profiling of mitochondrial-enriched material, a gel lane was cut into 60 gel slices (S1A Fig), excluding the very top and bottom sections of the gel. These sections were processed separately as they may contain aggregated proteins that would confound complexome analysis. Each slice was further diced into small cubes to aid trypsin digestion.

#### Blue Native (BN) Page for whole-cell samples

Two identical gels were run. For each of the two, 16.5 μL of cleared lysate containing 80 μg of protein was mixed with 1.5 μL loading dye (5 mM ACA, 5% (w/v) Coomassie brilliant blue (CBB) R-250) and 2 μL 50% (v/v) glycerol and separated by BN PAGE in NativePAGE 3%-12% Bis-Tris gels. The electrophoresis was performed in 200 mL of Blue Cathode Buffer (15 mM Bis-Tris/HCl pH 7.0, 50 mM Tricine, 0.002% (w/v) CBB R-250) and 150 mL Anode Buffer (50 mM Bis-Tris/HCl pH 7.0) under the following running conditions: a first step was performed at 80V and 15 mA for 10 min, followed by a second one at 100V and 20 mA for 3 hrs and 30 min, and finally a third one of additional 30 min at 130 V and 25mA. NativeMark was run along with the sample to estimate molecular weight sizes. Gels were both fixed in 25% (v/v) methanol and 10% (v/v) acetic acid on a shaker for 1 hr, at room temperature, and rinsed multiple times in MilliQ water.

For complexome profiling of whole-cell material, a gel lane was excised into 48 1.5 mm slices (S1A Fig), and these were transferred into a 96-well filter plate for further processing.

### In-gel trypsin digest

#### Mitochondrial samples

In-gel tryptic digestion was carried out on each gel slice. Briefly, diced gel slices were washed in 100 mM ammonium bicarbonate, followed by a wash in 50% acetonitrile and 100 mM ammonium bicarbonate. Cysteine reduction and alkylation were achieved *via* incubation in a solution of 45 mM DTT and 100 mM ammonium bicarbonate at 60 °C, followed by incubation in the dark with 100 mM iodoacetamide. Slices were washed again in 50% acetonitrile and 100 mM ammonium bicarbonate, followed by incubation in 100% acetonitrile. Trypsin digest was then performed overnight at 37 °C. Liquid was then removed from the gel slices and into to a 96-well microplate. The remaining gel slices were washed by incubation in 5% formic acid before the addition of acetonitrile, and the liquid transferred and pooled in the corresponding well of the microplate. Samples were then dried in a vacuum centrifuge, ready for MS analysis.

#### Whole-cell samples

Prior to the LC-MS analysis, protein digestion was performed on each gel slice. Briefly, gels were cut into smaller pieces and bleached with mixture of 50 mM ammonium bicarbonate (ABC) with acetonitrile (1:1 v/v). Gel was afterwards dried by pure acetonitrile and reduced with 10 mM DTT (dithiothreitol) in 100 mM ABC for 30 min at 60 °C. 55 mM chloroacetoamide in 100 mM ABC was added to the gel, incubated for 10 min at the room temperature and the gel was then washed twice with acetonitrile. Dry gel was mixed with trypsin that was resuspended in acetic acid, ABC and acetonitrile and incubated overnight at 37 °C. After digestion samples were mixed with 0.1% TFA in 60% acetonitrile and peptides were desalted using in-house made stage tips packed with C18 disks (Empore) according to [74].

### Mass spectrometry

#### Mitochondrial samples

Dried peptide residues were solubilized in 20 µL 5 % acetonitrile with 0.5 % formic acid using the auto-sampler of a nanoflow uHPLC system (Thermo Scientific RSLCnano). Online detection of peptide ions was by electrospray ionisation (ESI) mass spectrometry MS/MS with an Orbitrap Fusion MS (Thermo Scientific). Ionisation of LC eluent was performed by interfacing the LC coupling device to a NanoMate Triversa (Advion Bioscience) with an electrospray voltage of 1.7 kV. An injection volume of 5 µL of the reconstituted protein digest were desalted and concentrated for 10 min on trap column (0.3 × 5 mm) using a flow rate of 25 µl / min with 1 % acetonitrile with 0.1 % formic acid. Peptide separation was performed on a Pepmap C18 reversed-phase column (50 cm × 75 µm, particle size 3 µm, pore size 100 Å, Thermo Scientific) using a solvent gradient at a fixed solvent flow rate of 0.3 µl / min for the analytical column. The solvent composition was A) 0.1 % formic acid in water, B) 0.08 % formic acid in 80% acetonitrile 20% water. The solvent gradient was 4 % B for 1.5 min, 4 to 60 % for 50.5 min, 60 to 99 % for 3 min, held at 99 % for 7 min. A further 13 minutes at initial conditions for column re-equilibration was used before the next injection. The Orbitrap Fusion acquired full-scan MS in the range 310 to 2000 *m/z* for a high-resolution precursor scan at 120,000 RP (at 200 *m/z*), while simultaneously acquiring up to the top 15 precursors were isolated at 0.7 *m/z* width and subjected to CID fragmentation (35 % NCE) in the linear ion trap using rapid scan mode. Singly charged ions are excluded from selection, while selected precursors are added to a dynamic exclusion list for 30 sec.

#### Whole-cell samples

For LC/MS analysis, a Nano Reversed phase column (Ion Opticks, Aurora Ultimate XT C18 UHPLC column, 25 cm × 75 μm ID, 1.7 μm particles, 120 Å pore size, Ion Optics, Australia) was used for LC/MS analysis. Mobile phase buffer A was composed of water and 0.1% formic acid. Mobile phase B was composed of acetonitrile and 0.1% formic acid. Samples were loaded onto the trap column (C18 PepMap100, 5 μm particle size, 300 μm x 5 mm, Thermo Scientific) for 4 min at 18 μl/min loading buffer was composed of water, 2% acetonitrile and 0.1% trifluoroacetic acid. Peptides were eluted with mobile phase B gradient from 4% to 35% B in 15 min. Eluting peptide cations were converted to gas-phase ions by electrospray ionization and analysed on a Thermo Orbitrap Fusion (Thermo Scientific) by data dependent approach. Survey scans of peptide precursors from 350 to 1,400 m/z were performed in orbitrap at 120K resolution (at 200 m/z) with a 100 % ion count target. Tandem MS was performed by isolation at 1.6 Da with the quadrupole, HCD fragmentation with normalized collision energy of 30 % and 10 ms activation time. Fragmentation spectra were acquired in ion trap with scan rate set to Rapid. The MS2 ion count target was set to 150 % and the max injection time was 100 ms. Only those precursors with charge state 2–6 were sampled for MS2. The dynamic exclusion duration was set to 30 s with a 10-ppm tolerance around the selected precursor and its isotopes. Monoisotopic precursor selection was turned on. Cycle time was set to 1.5 s.

### Complexome profiling

Complexome profiling was performed as described in [32,33]. Briefly, RAW mass spectrometry data files for each individual gel slice were analysed using MaxQuant (v2.0.3.0 or v2.4.13.0) [75], accessed using the Galaxy server [76], using default settings. For protein identification, peptides were searched against a custom database containing the *P. marinus* proteome (Uniprot UP000007800) and protein sequences for mitochondrial encoded proteins CoxI, CoxIII and CytB [25]. Protein abundance values were calculated using the iBAQ method and profiles analysed using Nova software (v0.5.8) [77]. Profiles were normalised using the maximum normalisation function and hierarchical clustering performed with the Pearson Correlation distance function. Heatmaps were visualised using GraphPad Prism software (v10.6.1).

### Phylogenetic analysis

The ATPTG1 protein sequence from *P. marinus* (Pmar_PMAR013021) was used as a query in BLASTp search against the dataset of 132 alveolate species from EukProt database v.3[78] (S7 Table) with an *e*-value threshold of 1*e*-5. Retrieved sequences were aligned using MAFFT v.7.505 with the L-INS-i algorithm [79] and trimmed using trimAl v1.4.rev15 [80] with the -gt 0.8 option. Phylogenomic analysis was performed using IQ-TREE v.3.0.1 [81] with automatic model selection and branch support values estimated using Shimodaira–Hasegawa-like aLRT test. The resulting tree was visualized with Interactive Tree Of Life (iTOL) web server [82].

### Immunofluorescence assay in *Toxoplasma gondii*

*Toxoplasma gondii* was cultured as described previously [10]. The sequence of TGGT1_224932 was synthesised with the addition of a 5’ *EcoRI* restriction site and expression sequence, a 3’ *NsiI* restriction site, as well as the remove of internal *NsiI* sites. This sequence was then inserted into the pTUB8mycGFPMyoATy expression vector [83]. The resulting plasmid was used to transfect *Toxoplasma* which were then inoculated onto HFFs grown on glass coverslips. Cells were fixed with 4% paraformaldehyde after 1 day. An immunofluorescence assay was performed as described previously [10], using primary antibodies against Ty (1:800, anti-mouse,[84]) and TgTom40 (1:1,000, anti-rabbit,[42]), and imaged on a Nikon inverted AX/ AX-R Laser-Scanning Confocal Microscope using 63x objective. Z-stacking was performed on Nikon’s NIS-Elements software using high-speed resonant scanner. The images were analysed using FIJI ImageJ X-64 software [85].

## Acknowledgements

We would like to thank Hannah Bridges and Judy Hirst (University of Cambridge) for the gift of bovine mitochondrial membrane. The help of Anzhelika Butenko (Institute of Parasitology, Biology Centre) with the phylogenetic analysis of ATPTG1 proteins is appreciated. We are grateful to Shikha Shikha (University of Glasgow) for discussion of the mitoribosomal subunits analysis. The authors gratefully acknowledge the Mass Spectrometry Facility, part of the MVLS Shared Research Facility and Laboratory of Mass Spectrometry at Biocev research center, Faculty of Science, Charles University, Prague, for their support & assistance in this work. This work was supported by Sir Henry Wellcome Fellowship (grant number 221681/Z/20/Z), Wellcome ISSF feasibility award (grant number 204820/Z/16/Z) and a Medical Research Council Career Development Award UKRI325 (grant reference: UKRI/MR/B000325/1) (to A. E. M); by the Czech Science Foundation grant 23-06479X (to J.L.) and by Wellcome Trust’s Wellcome Investigator Award - 217173/Z/19/Z, and Wellcome Discovery Award - 310879/Z/24/Z (to L.S.).

---

**Figure S1:**
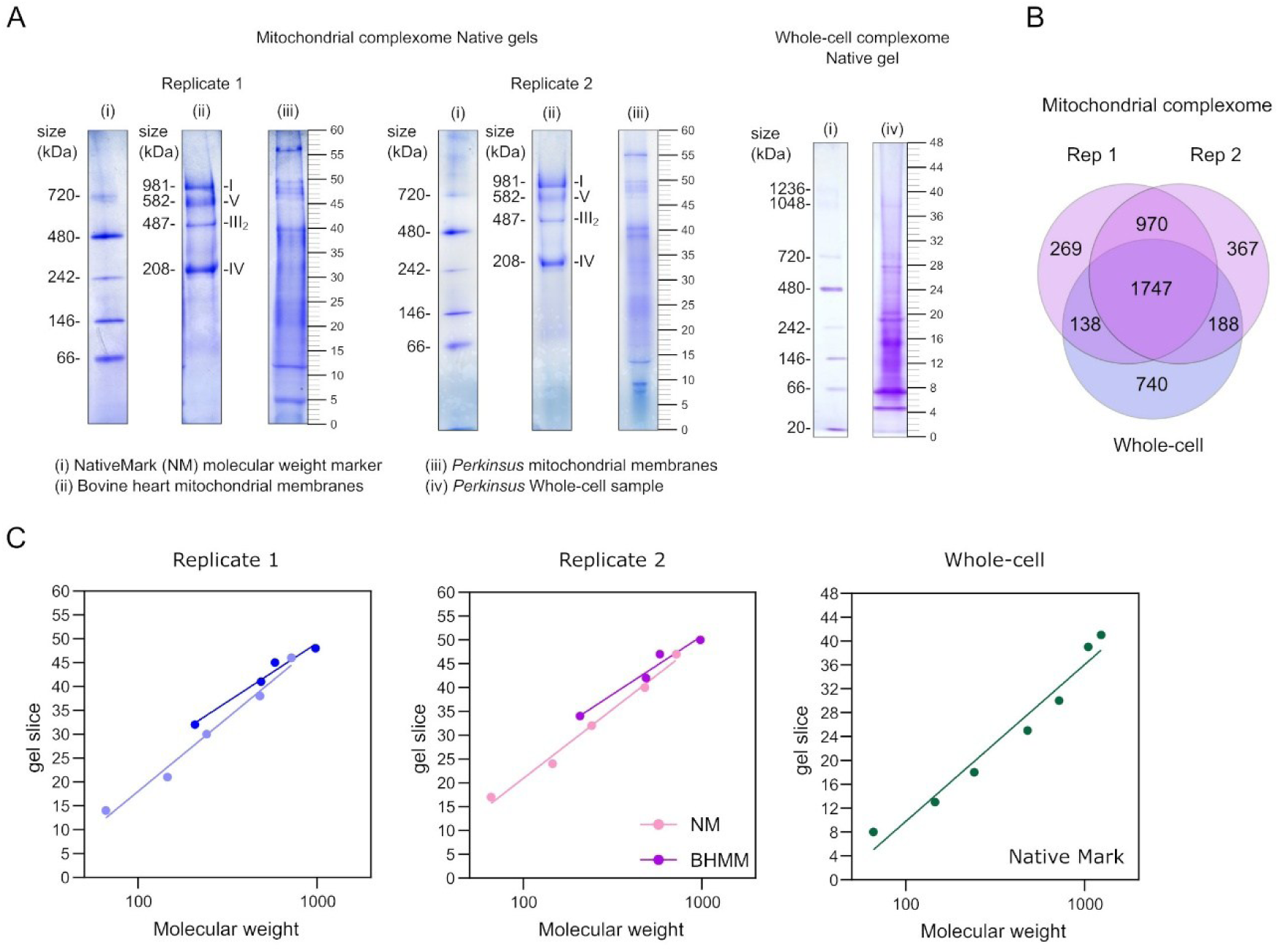
Generating complexome profiles. (A) Coomassie-stained Native PAGE gels showing molecular weight markers (i-NativeMark, ii-Bovine mitochondrial membranes), *Perkinsus* mitochondrially-enriched material (iii) and *Perkinsus* whole-cell material (iv) used in generating the complexome. Molecular weight markers and *Perkinsus* material were run on the same gel and aligned, with corresponding gel slices numbers (1-60) shown on the right-hand side. For the mitochondrially-enriched material, gels from both replicate 1 and 2 are shown. (B) Comparison of proteins identified in each repetition of the mitochondrial and the whole-cell samples. (C) Mass calibration analysis using NativeMark and Bovine mitochondrial membrane molecular weight markers. Molecular weight and corresponding gel slice were plotted and a non-linear regression, semi-log line plotted. Full details of mass calibration are in Table S3.

**Figure S2:**
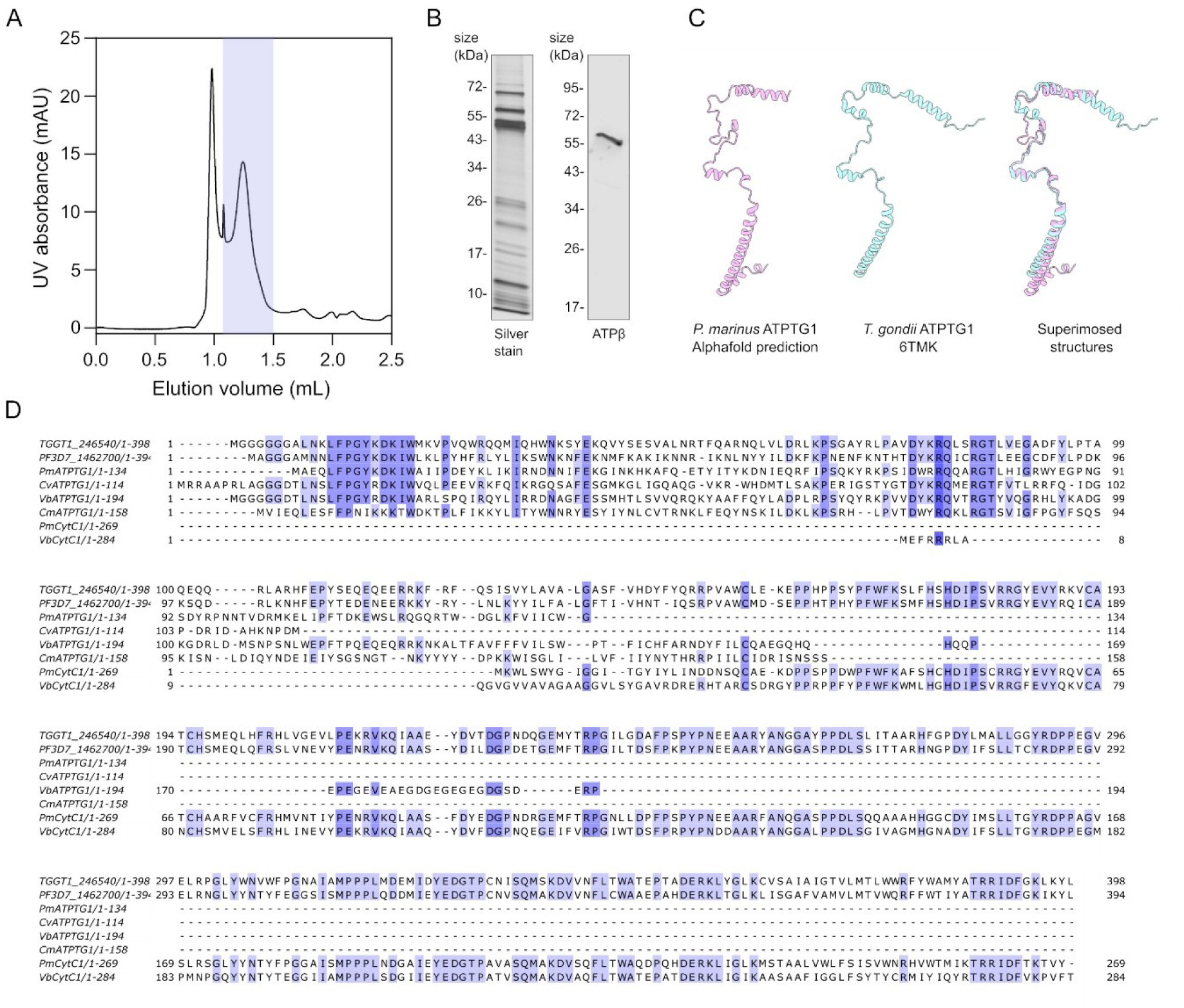
Analysis of ATP synthase subunit ATPTG1. (A) Chromatogram displaying UV 1_280 of gel filtration of purified ATP synthase using a Superose 6 increase 3.2/300 column. Left-most peak represents void-volume. The fractions highlighted in blue were combined and concentrated using a 100 MWCO spin concentrator before analysis via immunoblot and mass spectrometry. (B) SDS-PAGE analysis of purified ATP synthase. Gel was either silver stained to visualize protein subunits or subjected to immunoblot analysis with the ATPβ antibody. (C) Structural prediction of *P. marinus* ATPTG1 (predicted using Alphafold3 [87]) (left), the structure of *T. gondii* ATPTG1 (PDB 6TMK; [17]) (center) and the superimposition of the two structures (RMSD = 1.122 Å) (right). Visualized using ChimeraX [86]. (D) Alignment of ATPTG1, CytC1 and fusion proteins from across the myzozoa. ATPTG1 proteins: *Perkinsus marinus* Pmar_PMAR013021; *Chromera velia* Cvel_13300*; Vitrella brassicaformis Vbra_20035*; *Cryptosporidium muris* CMU_009920. CytC1 proteins: *Perkinsus marinus* Pmar_PMAR028703; *Vitrella brassicaformis* Vbra_21749. Fusion proteins: TGGT1_246540 from *Toxoplasma gondii*; PF3D7_1462700 from *Plasmodium falciparum*. Alignments were made using Clustal Omega and visualized using JalView (v10.0.7) [88]. Colour coding depicts percent identity.

**Figure S3:**
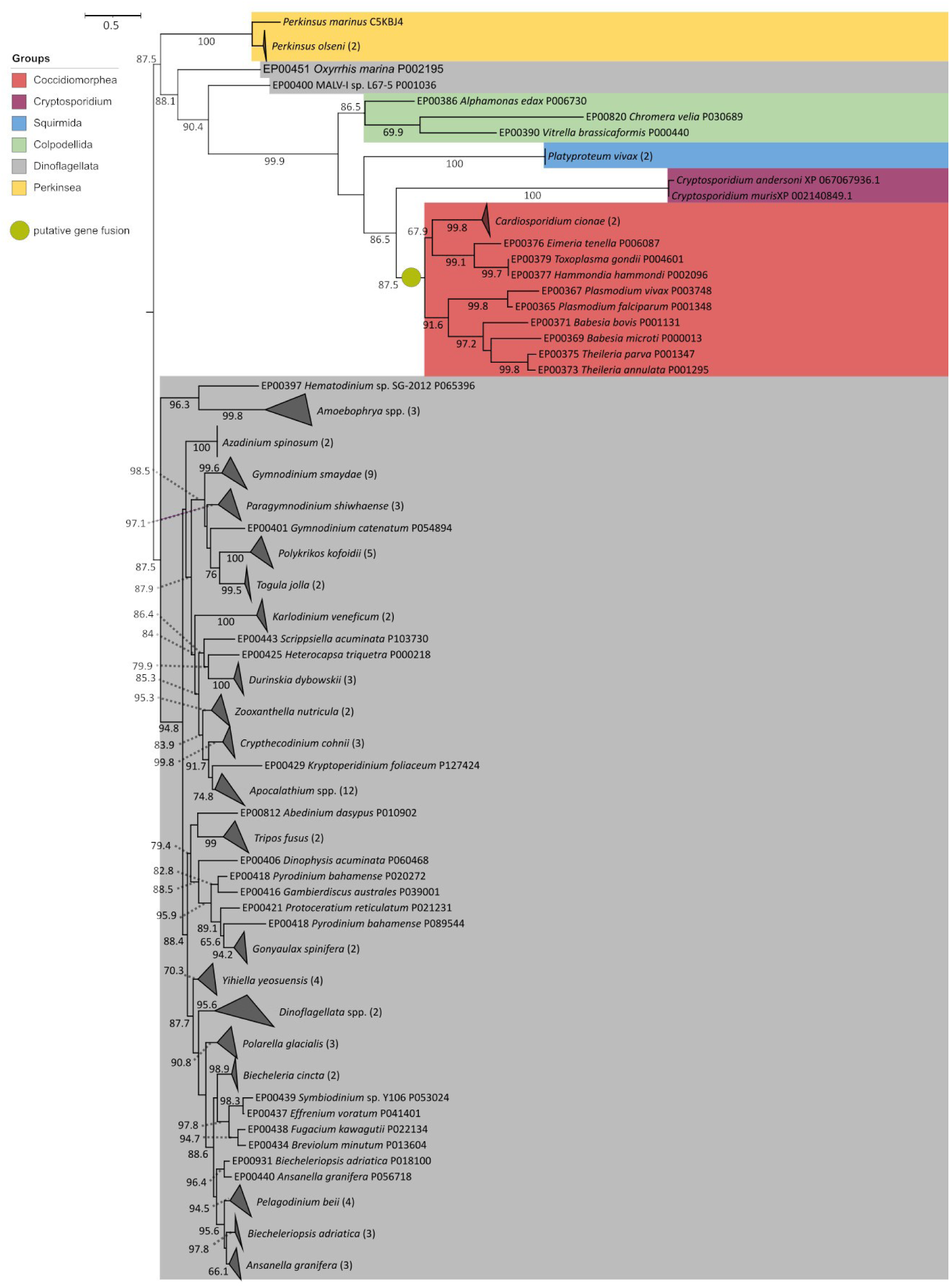
Phylogeny of ATPTG1 homologues from various myzozoans. A maximum-likelihood phylogenetic tree inferred with IQ-TREE v.3.0.1 on the trimmed protein alignment containing 170 amino acid positions. The tree was inferred with the LG+I+G4 model automatically selected according to Bayesian information criterion. Branch support values represent Shimodaira–Hasegawa-like aLRT (SH-aLRT) test scores; only values ≥60 are shown. The horizontal bar indicates number of substitutions per site.

## Notes

### Competing Interest Statement

The authors have declared no competing interest.

### Summary of Updates

Added italics to submission title. Perkinus marinus was not previously italicised- it has been updated accordingly.

